# FKSFold: Improving AlphaFold3-Type Predictions of Molecular Glue–Induced Ternary Complexes with Feynman–Kac–Steered Diffusion

**DOI:** 10.1101/2025.05.03.651455

**Authors:** Jian Shen, Shengmin Zhou, Xing Che

## Abstract

We introduce an AI model, FKSFold, that uses Feynman–Kac steered diffusion as an inference-time strategy to improve AlphaFold3-type predictions of molecular-glue induced ternary complexes. FKSFold augments reverse diffusion with a Feynman–Kac derived steering term and uses the interface predicted TM-score (ipTM) as the guiding potential, coupled with particle-based sampling and adaptive resampling to bias trajectories toward high-quality interfaces without retraining the base models. Implemented on Chai-1r and Boltz-2 (named FKSFold-Chai and FKSFold-Boltz) – two open-source AF3-type architectures – FKSFold was benchmarked on eight ternary systems. FKSFold-Chai recovered three challenging complexes with sub-3 Å accuracy: VHL:MG:CDO1 (DockQ 0.922, iRMSD 0.629, fnat 0.963), FKBP12:MG:mTOR-FRB (DockQ 0.590, iRMSD 1.813, fnat 0.519), and FKBP12:MG:BRD9 (DockQ 0.841, iRMSD 0.801, fnat 0.913), while unmodified baselines failed. Other cases highlighted challenges – flexible loop rearrangements (e.g., CRBN:MG:VAV1-SH3c) and large conformational search spaces (e.g., NEK7, QDPR, HDAC1) – and, empirically, using few particles, short resampling intervals, and moderate-to-high lambda (steering strength) gave the best exploration–exploitation balance. Although the strategy is theoretically model-agnostic, on Boltz-2 ipTM head instability currently limits observed gains. The code for the two implementations, FKSFold-Chai and FKSFold-Boltz, is available at https://github.com/YDS-Pharmatech/FKSFold-Chai and https://github.com/YDS-Pharmatech/FKSFold-Boltz.

## Introduction

The prediction of protein-protein interactions induced by small molecules, known as molecular glues, represents a frontier challenge in computational chemistry.^1,2^ Molecular glues are small monofunctional compounds that mediate protein-protein interactions. They function through two main mechanisms. Degradative molecular glues typically bind E3 ubiquitin ligases to recruit target proteins for ubiquitination and subsequent proteasomal degradation.^3,4^ Non-degradative molecular glues stabilize native protein interactions or induce new functional protein complexes without causing degradation. ^5^ The therapeutic value of these compounds is exceptionally high due to their catalytic mechanism of action, enabling them to function at lower doses with greater efficiency than traditional inhibitors,^6^ their ability to target proteins previously considered undruggable due to lack of defined binding pockets,^6,7^ and their favorable drug-like properties including lower molecular weight and improved cell permeability compared to other targeted protein degradation approaches.^3^ These molecular glues enable novel therapeutic strategies for previously undruggable targets such as transcription factors IKZF1 and IKZF3 in multiple myeloma, ^8^ oncogenic *β*-catenin,^9^ p53,^10^ and scaffold proteins lacking defined binding pockets.^6^

Despite their therapeutic potential, the discovery of new molecular glues has been largely serendipitous, hindering broad translational efforts in this promising field.^2,11^ The molecular glue community is actively pursuing a more rational design approach for previously undruggable targets.^12^ Computational methods offer promising solutions in this pursuit. Accurate ternary structure prediction would enable more efficient screening of hits and optimization of lead compounds, guiding medicinal chemistry efforts and reducing reliance on resource-intensive experimental methods.^13,14^

Recent advances in computational structural biology offer promising tools to address these challenges. At the forefront, AlphaFold 3, ^15^ developed by Google DeepMind and Isomorphic Labs, represents a significant breakthrough in biomolecular interaction complex prediction with its novel architecture that combines a Pairformer module and a diffusion-based model. At the core of AlphaFold 3 is its ability to replace the previous Evoformer with the Pairformer for generating intermolecular interaction hypotheses, then employing a diffusion module that directly predicts raw atom coordinates through iterative denoising on the structural sketches. While AlphaFold 3 achieves unprecedented accuracy for many protein-protein interactions,^16^ accurately modeling molecular glue-induced ternary complexes remains difficult due to the complex energetic landscape that governs the formation of these three-body systems. ^1,14^

In this paper, we disclose technical details of an early approach we explored for molecular glue-induced ternary complex structure prediction using Feynman-Kac (FK) steering in the diffusion modules of AlphaFold3-type models. Steering diffusion represents a straightforward approach to adjust diffusion sampling, a strategy later formalized and published under the name “FK Steering.”^17–20^ We adopt this terminology throughout our paper. To evaluate the generalizability of our approach, we implemented FK steering modifications to both Chai-1r and Boltz-2 architectures, with Chai-1r demonstrating consistently superior performance in our test cases. While we recently reported the successful results of our YDS-GlueFold model^21^ in predicting 8 novel molecular glue-induced ternary structures validated by experimental data, here we share insights from our earlier FK steering experiments which preceded YDS-GlueFold’s development. Our approach leverages the mathematical framework of Feynman-Kac formalism to modify the diffusion process during inference, using interface predicted TM-score (ipTM) as a reward function to guide sampling toward high-quality protein-protein interfaces in ternary complexes.

Despite mixed overall results, our initial experiments with FK steering demonstrated promising potential in several key test cases. FK steering successfully predicted 3 out of the 8 cases (VHL:MG:CDO1, FKBP12:MG:mTOR-FRB, and FKBP12:MG:BRD9) that were later solved by YDS-GlueFold, with RMSD values below 3Å. The remaining cases presented greater challenges, very likely because those proteins containing flexible regions or large conformational sampling spaces. Our hyperparameter tuning studies revealed the sensitivity of the FK steering approach to various parameters, including particle numbers, diffusion path length, temperature parameters, potential function design, and sigma threshold values. By documenting this early approach, we provide insights into the evolution of our technical methodology toward molecular glue-induced complex prediction. The FK steering approach represents a theoretically grounded strategy for incorporating domain-specific knowledge directly into the diffusion process without requiring additional training. While it achieved only partial success in our initial implementation, the lessons learned from these experiments were instrumental in developing the more comprehensive and successful YDS-GlueFold model. This work contributes to the expanding toolkit of computational methods for addressing the complex challenge of modeling novel complex interactions.

## Methods

### Diffusion Models in Protein Complex Prediction

Diffusion models have emerged as powerful generative models across various domains, including protein structure prediction and complex modeling. These models work by defining a forward process that gradually adds noise to data and a reverse process that reconstructs data from noise.^22,23^ In the context of molecular glue-induced ternary complex prediction, the forward process can be formulated as:

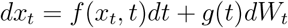

where *t* = 0 represents the original data and *t* = 1 represents the fully noisy state. *x*_*t*_ represents the ternary complex structure (comprising two proteins and a molecular glue) at diffusion time *t, g*(*t*) relates to the variance schedule, which can be derived from the relationship:

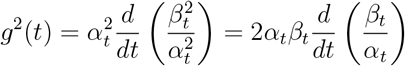

where *β*_*t*_ represents the noise variance at time *t*. And *W*_*t*_ is a standard Brownian motion. *f* (*x*_*t*_, *t*) is the drift term, specifically:

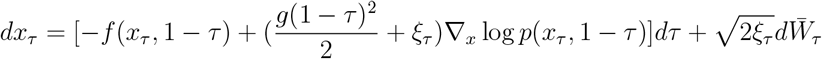

where *α*_*t*_ controls the decay of the original signal.

Under this formulation, the conditional distribution *p*(*x*_*t*_|*x*_0_) follows a Gaussian distribution 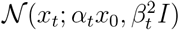, which allows for direct sampling from any arbitrary timestep.

The reverse process aims to recover the ternary complex structure by solving:

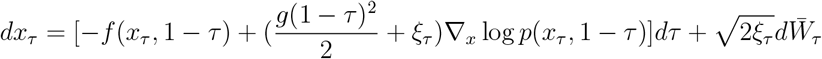

where *τ* = 1 − *t, ∇*_*x*_ log *p*(*x*_*τ*_, 1 − *τ*) is the score function representing the gradient of the log probability density, and 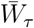 is a standard Brownian motion in the reverse time direction. This formulation follows from Anderson’s theorem on the time-reversal of diffusion processes, which states that if a forward process follows an SDE with drift *f* (*x*_*t*_, *t*) and diffusion coefficient *g*(*t*), then its time-reversal follows an SDE with an additional drift term involving the score function ∇_*x*_ log *p*(*x*_*τ*_, 1 −*τ*), where *p*(*x*_*τ*_, 1 −*τ*) is the marginal probability density at time *t*.

The framework allows flexibility in the reverse process by introducing a parameter *ξ*_*τ*_ .As demonstrated by Song et al., as long as 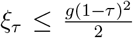, the marginal distributions of the reverse process remain consistent with the forward process. This important insight reveals that different reverse processes with varying noise levels can generate samples from the same distribution. When 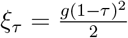, we recover the standard reverse SDE, while *ξ*_τ_ = 0 yields a deterministic ODE formulation:

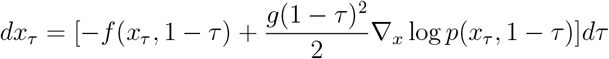

This deterministic approach, analogous to the Probability Flow ODE, enables faster sampling without introducing additional noise, significantly accelerating inference while maintaining generation quality. This concept generalizes the Denoising Diffusion Implicit Models (DDIM)^24^ approach to continuous time diffusion models.

In AlphaFold3-type models applied to ternary complexes, diffusion processes operate on structural representations that must simultaneously capture protein-protein interfaces and protein-ligand interactions. This includes relative orientations between proteins, interface residue conformations, and small molecule positioning at the interface region, making the prediction task substantially more complex than individual protein structure prediction.

### Feynman-Kac Formalism

The behavior of the diffusion model can be adjusted through multiple approaches, such as simply increasing diffusion steps or modifying the diffusion noise scheduler to achieve an optimal balance between stability and diversity. These parameter adjustments, while seemingly incremental, can impact the quality of structural predictions by altering the exploration-exploitation trade-off inherent in diffusion processes. However, these approaches are unable to guide the diffusion process toward enhanced intermolecular interactions. Such basic modifications merely change the sampling dynamics uniformly across the entire structural space, without specifically directing the model toward regions of higher biological relevance. More sophisticated approaches like classifier guidance have been proposed to steer diffusion models toward desired attributes, but these typically require additional training or fine-tuning of separate classifier networks.

What is needed is a mathematically rigorous framework that can incorporate domain-specific knowledge about protein interfaces while maintaining the theoretical foundations of the diffusion process. Such a framework should be able to bias sampling toward biophysically plausible conformations without requiring extensive retraining of the underlying model or compromising the diversity of generated structures. The Feynman-Kac formalism provides precisely such a framework, offering a principled way to integrate reward functions like interface quality metrics directly into the diffusion dynamics.^25,26^

The Feynman-Kac formula establishes a connection between partial differential equations and stochastic processes. The steering term *h*(*x*_*τ*_, 1 − *τ*) can be defined explicitly in terms of the potential function *V* (*x*_*τ*_, 1 − *τ*) as follows:

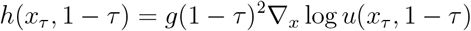

Where *u*(*x*_*τ*_, 1 − *τ*) is the solution to the Feynman-Kac PDE:

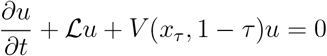

With the final condition *u*(*T, x*) = *ϕ*(*x*), where ℒ is the generator of the diffusion process and *V* is a potential function, the solution of *u*(*x*_*τ*_, 1 − *τ*) can be expressed as

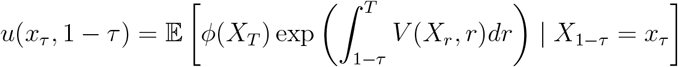

Where we denote the stochastic process by uppercase *X*_*t*_ and its specific state at time *t* by lowercase *x*_*t*_. In the Feynman-Kac formulation, the condition *X*_1−*τ*_ = *x*_*τ*_ indicates that the stochastic process is fixed in the known state *x*_*τ*_ at time 1 − *τ* . This establishes that the steering term *h* is derived from the gradient of the logarithm of the solution to the FK PDE, which incorporates the potential function *V* .

### FK Steering for Diffusion Models with ipTM Guidance

We propose a novel approach that integrates FK formalism with diffusion models for molecular glue-induced ternary complex prediction. The key idea is to modify the reverse diffusion process by introducing a steering term derived from the FK formula, guided by ipTM scores. The modified reverse process becomes:

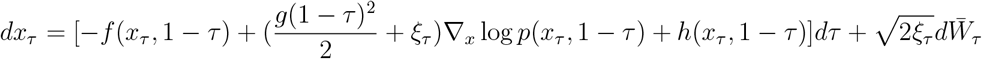

where *h*(*x*_*τ*_, 1 − *τ*) is the steering term derived from the FK formula: *h*(*x*_*τ*_, 1 − *τ*) = *g*(1 − *τ*)^2^∇_*x*_ log *u*(*x*_*τ*_, 1 − *τ*)

Here, *u*(*x*_*τ*_, 1 −*τ*) is the solution to the FK PDE incorporating domain-specific knowledge about protein-protein interfaces and molecular glue binding through the potential function *V* (*x*_*τ*_, 1 − *τ*).

Similar to how the probability flow ODE provides a deterministic alternative to SDE sampling in diffusion models, our framework enables more directed sampling toward highquality interfaces while maintaining the theoretical guarantees of the underlying diffusion process.

Unlike previous approaches that apply gradient guidance uniformly throughout the generation process, our FK steering framework adaptively focuses computational resources on the most promising trajectories through resampling. Furthermore, in contrast to methods that rely solely on physical scoring functions, our approach leverages the ipTM neural network predictor that captures complex patterns in protein-protein interfaces learned from structural data.

A key challenge in FK steering is maintaining particle diversity while biasing toward high-quality structures. We address this by employing adaptive resampling based on effective sample size and by tuning the temperature parameter *λ* in our potential function. Lower values of *λ* promote exploration of diverse binding modes, while higher values focus sampling on the highest ipTM-scoring conformations.

#### Lambda Weight Parameter

The lambda weight (*λ*) functions as a temperature parameter in our resampling mechanism. Mathematically, this parameter scales the reward values (ipTM scores based) when calculating sampling probabilities:

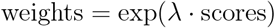

### Potential Function Design with ipTM Integration

The effectiveness of FK steering largely depends on the design of the potential function *V* (*t, x*). We propose a potential function that incorporates domain knowledge about protein-protein-ligand interactions, with a central focus on the interface quality:

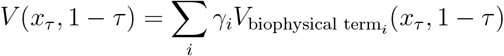

where 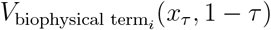 could be:

- *V*_*ipTM*_ is based on the predicted interface quality using the ipTM score, guiding the diffusion process toward conformations with higher-quality protein-protein interfaces. This term directly captures the likelihood of forming stable protein-protein contacts in the presence of the molecular glue.
- *V*_*phys*_ captures physical constraints such as steric clashes, hydrogen bonding patterns, and electrostatic interactions at the interface.
- *V*_*chem*_ incorporates chemical knowledge about molecular glue binding, including pharmacophore features and key interaction motifs.
- *V*_*prior*_ encodes prior knowledge about similar ternary complexes and common molecular glue binding modes.

The coefficients *γ*_*i*_ control the relative importance of each term and can be adjusted based on the confidence in each information source, with *γ* for *V*_*ipTM*_ typically assigned a higher weight to emphasize the importance of interface quality. In practice, we compute the gradient term ∇_*x*_ log *u*(*x*_*τ*_, 1 − *τ*) using automatic differentiation of our potential function. The ipTM score is evaluated on intermediate denoised states.

#### Potential Function Type

The potential function type determines how reward signals are integrated into the FK steering process. We implemented three variants:

- **Vanilla potential**: Uses current scores directly: exp(*λ ·* current scores)
- **Difference potential**: Emphasizes improvement over historical scores: exp(*λ·*(current scores− historical scores))
- **Maximum potential**: Considers the maximum between current and historical scores: exp(*λ ·* max(current scores, historical scores))

### Implementation in the Diffusion Module for Ternary Complex Prediction

Our implementation integrates FK steering within the diffusion module of AlphaFold3-type models while maintaining the core structure module intact. We focus on modifying the inference process to enhance sampling for molecular glue-induced ternary complexes through a particle-based approach that leverages the ipTM reward function.

Our objective is to guide the diffusion process toward enhanced intermolecular interactions. Therefore, we selected the Interface Predicted TM-score (ipTM) as our main quality metric for evaluating each diffusion step. The efficacy of ipTM stems from its focused assessment of interface regions—critical determinants of protein-protein interaction specificity and affinity. Additionally, ipTM provides a quantitative standard enabling comparative reliability assessments across different complex predictions. Importantly, we can directly leverage AlphaFold3’s confidence head output to estimate ipTM, adhering to our principle of avoiding post-training modifications.

For diffusion path optimization, we maintain *k* parallel diffusion paths and implement a comprehensive sampling strategy informed by Feynman-Kac formalism. At each diffusion step, we execute a standard forward pass through the model for all particles to obtain the score function ∇_*x*_ log *p*(*x*_*τ*_, 1 − *τ*). We then leverage the model’s confidence assessment networks to predict ipTM scores and other scores for each partially denoised structure, capturing the quality of protein-protein interfaces. Using these scores, we calculate FK steering terms that incorporate both interface quality metrics and additional biophysical potential functions. The diffusion update for each particle combines the standard score-based update with our FK steering term, guiding the sampling process toward conformations with superior interface characteristics. Crucially, we perform periodic resampling of particles using Multinomial sampling weighted by current ipTM scores, effectively concentrating computational resources on the most promising structural configurations while maintaining sufficient diversity. This approach elegantly balances exploration and exploitation without requiring any additional training of the underlying model, demonstrating versatility across various reward functions and potential adaptability to different generative modeling frameworks beyond protein structure prediction.

## Results

### Performance on Molecular Glue Complexes

In our early experiments with FK steering, we evaluated its performance on a set of 8 molecular glue-induced ternary complexes—the same set that was later successfully predicted by our YDS-GlueFold model. Table 1 summarizes the performance of FK steering in these cases:

**Table 1:** Performance of FKSFold on molecular glue-induced ternary complexes.

| Complexes | RMSD (Å) | Success | Chai-1r* | Boltz-2* | YDS-GlueFold* |
| --- | --- | --- | --- | --- | --- |
| VHL:MG:CDO1 | 1.42 | ✓ | × | × | ✓ |
| FKBP12:MG:mTOR-FRB | 2.14 | ✓ | × | × | ✓ |
| FKBP12:MG:BRD9 | 1.45 | ✓ | × | × | ✓ |
| CRBN:MG:mTOR-FRB | 8.79 | × | × | × | ✓ |
| CRBN:MG:NEK7 | 21.65 | × | × | × | ✓ |
| CRBN:MG:VAV1-SH3c | 6.65 | × | × | × | ✓ |
| FKBP12:MG:QDPR | 11.36 | × | × | × | ✓ |
| KBTBD4:MG:HDAC1 | 19.17 | × | × | × | ✓ |

\*Data from YDS-GlueFold: Surpassing AlphaFold 3-Type Models for Molecular Glue-Induced Ternary Complex Prediction.<sup>21</sup> A prediction was considered successful if the root-mean-square deviation (RMSD) was below 4 Å.

Our FK steering approach successfully predicted 3 out of the 8 cases (VHL:MG:CDO1, FKBP12:MG:mTOR-FRB and FKBP12:MG:BRD9) with RMSD below 3Å, which is generally considered the threshold for a successful prediction. RMSD calculations were performed using reference structures derived from either PDB entries or final predictions generated by our YDS-GlueFold model. While the method shows some promise, it struggled with the majority of cases, which later motivated our development of the more comprehensive YDS-GlueFold approach.

### Detailed Analysis of Successfully Predicted Cases

#### VHL:MG:CDO1: A Novel VHL-based Molecular Glue System

VHL:MG:CDO1 represents a particularly challenging VHL-based molecular glue degrader system with a complex binding geometry. Figure 2(a) shows a detailed comparison of FK steering prediction against the experimental structure.

**Figure 1:**
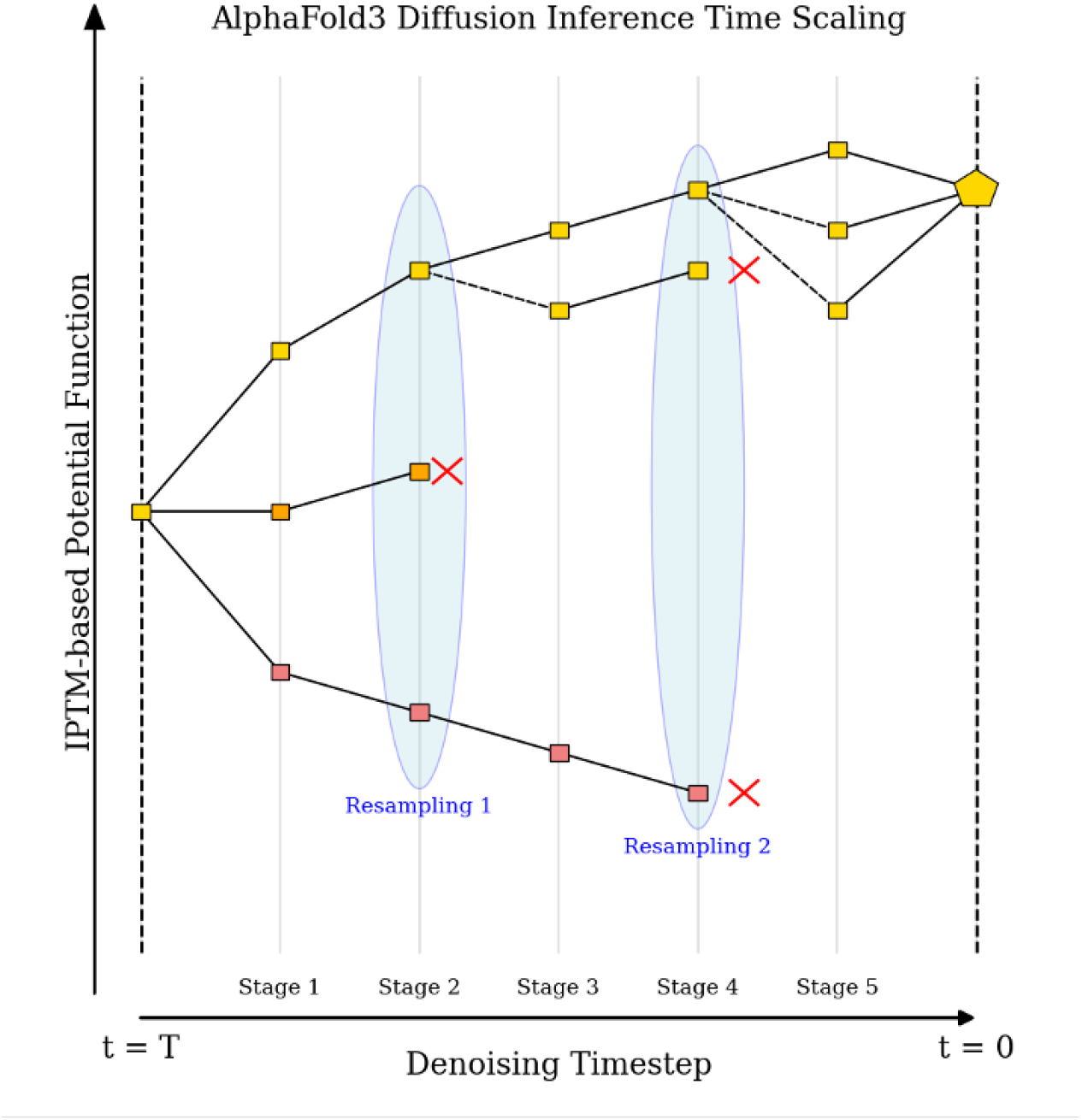
FK Steering Diffusion Inference Time Scaling using IPTM-based Potential Function

**Figure 2:**
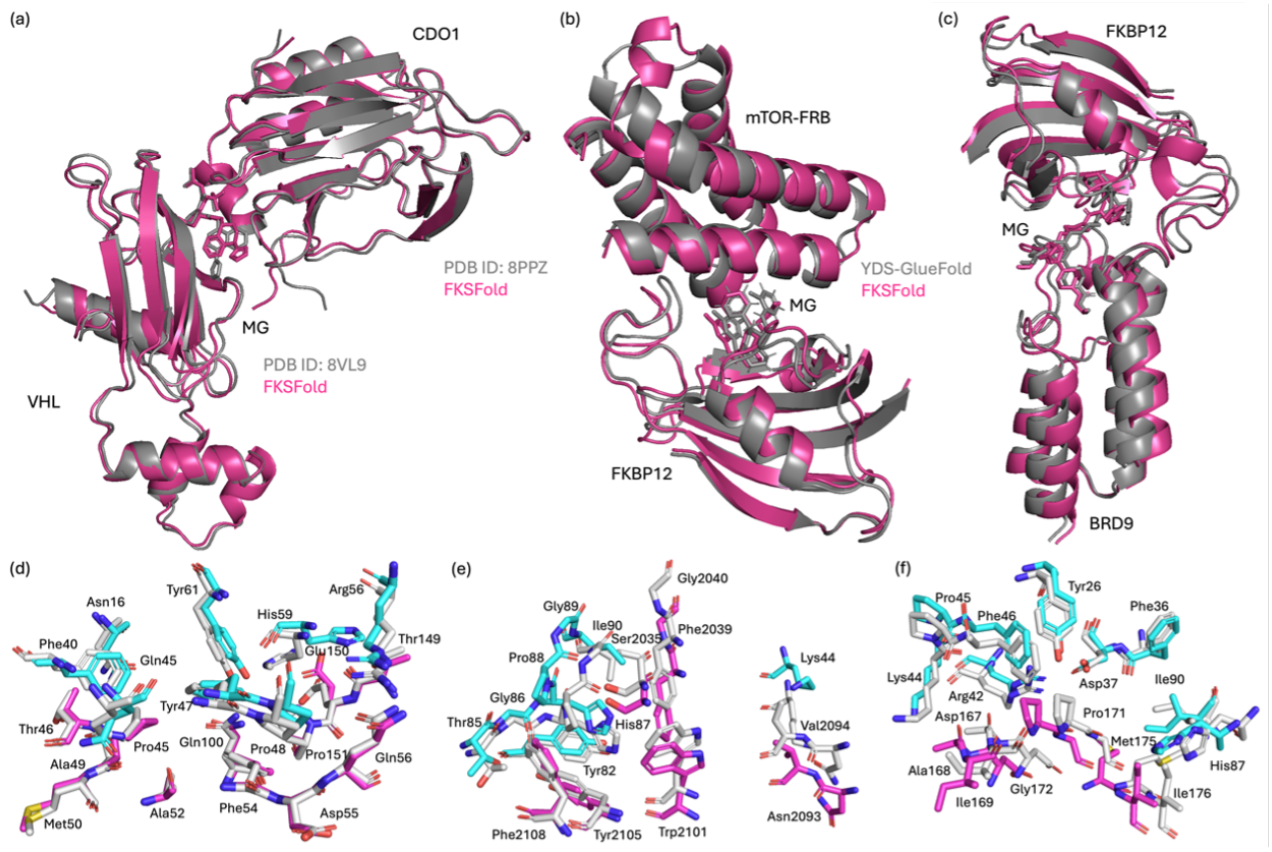
Structure predictions for the three successfully predicted cases, (a) VHL:MG:CDO1,(b) FKBP12:MG:mTOR-FRB and (c) FKBP12:MG:BRD9, using FK steering. The reference structures (either PDB structure of YDS-GlueFold structure) are shown in gray, with FK steering predictions in red cartoon representation. Key interface residues are highlighted in (d) VHL:MG:CDO1, (e) FKBP12:MG:mTOR-FRB and (f) FKBP12:MG:BRD9. Residues from FK Steering predicted VHL or FKBP12 are shown in cyan, residues from FK Steering predicted CDO1, mTOR-FRB or BRD9 are shown in magentas, and residues from reference structures are shown in grey.

The FK steering approach successfully captured the critical interactions at the interface (Figure 2(d)), particularly the precise positioning of the molecular glue that mediates the protein-protein interaction. The substantial improvement in RMSD compared to Chai-1 prediction demonstrates the effectiveness of the ipTM-guided steering approach for this complex case.

We subsequently conducted additional experiments to validate the reproducibility of the results presented in Figure 2 and to observe the actual diffusion trajectories.

As demonstrated in Figure 3, FK steering indeed contributes to improved diffusion performance in the molecular glue domain. To further validate the generalizability of our approach, we also evaluated FK steering on Boltz-2, which presents additional challenges due to architectural differences. Specifically, Boltz-2 employs a smaller confidence head that tends to produce higher but more unstable scores. Nevertheless, through FK steering, we achieved superior results compared to the baseline performance.

**Figure 3:**
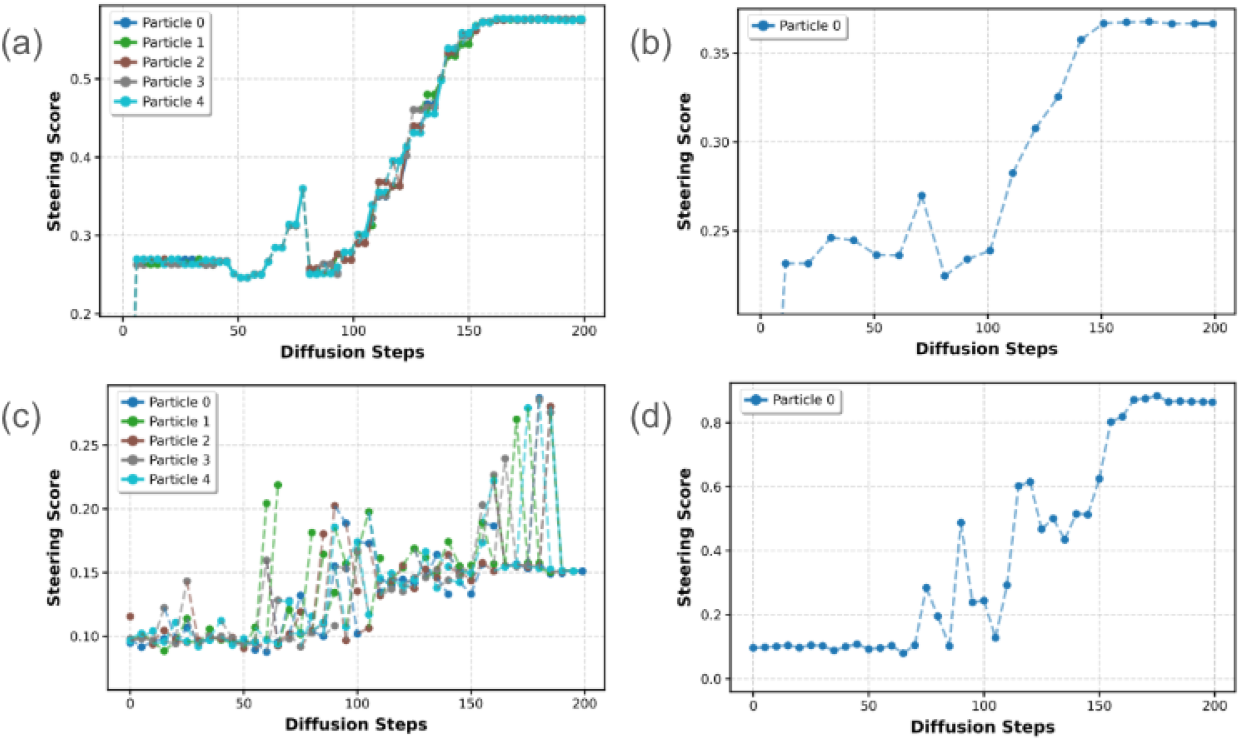
Internal diffusion score trajectories for the VHL:MG:CDO1, (a) Chai-1 with ipTM-guided steering and (b) Chai-1 default diffusion and (c) Boltz-2 with ipTM-guided steering and (d) Boltz-2 with default diffusion

The quantitative results in Table 2 demonstrate substantial improvements achieved through FK steering across multiple evaluation metrics. For the Chai baseline, FKSFold-Chai dramatically enhanced performance, with DockQ scores improving from 0.012 to 0.922, interface RMSD (iRMSD) decreasing from 15.899 Å to 0.629 Å. The fraction of native contacts (fnat) increased from 0.000 to 0.963, indicating successful recovery of critical intermolecular interactions.

**Table 2:**
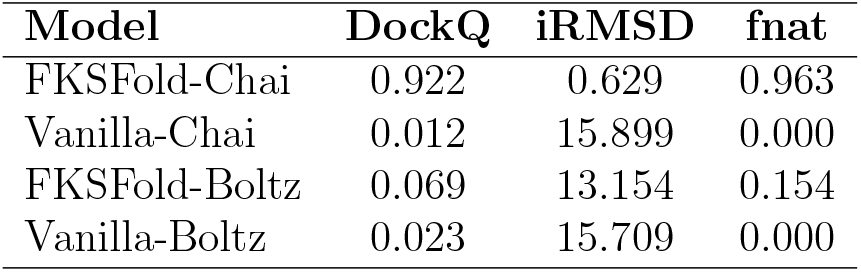
VHL:MG:CDO1 Performance comparison of FK steering across different models on molecular glue-induced ternary complex prediction.

**Table 3:** FKBP12:MG:mTOR-FRB Performance comparison of FK steering across different models on molecular glue-induced ternary complex prediction.

| Model | DockQ | iRMSD | f<sub>nat</sub> |
| --- | --- | --- | --- |
| FKSFold-Chai | 0.590 | 1.813 | 0.519 |
| Vanilla-Chai | 0.076 | 9.172 | 0.036 |
| FKSFold-Boltz | 0.151 | 5.420 | 0.080 |
| Vanilla-Boltz | 0.064 | 10.162 | 0.030 |

**Table 4:**
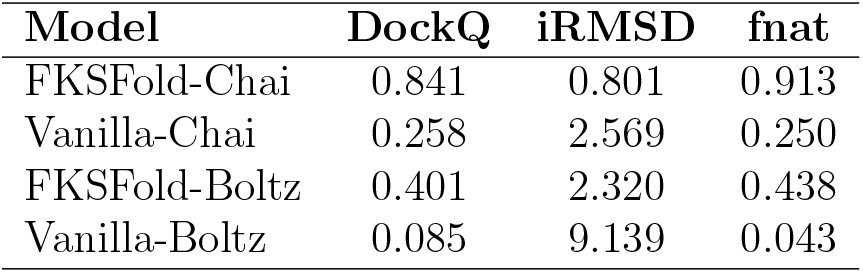
FKBP12:MG:BRD9 Performance comparison of FK steering across different models on molecular glue-induced ternary complex prediction.

#### FKBP12:MG:mTOR-FRB: A Challenging Non-E3 Ligase Molecular Glue System

FKBP12:MG:mTOR-FRB represents a distinct molecular glue system where FKBP12 functions as a prolyl isomerase rather than an E3 ligase. Figure 2(b) shows a detailed comparison of FK steering prediction against the experimental structure from PDB ID: 8PPZ. The FK steering approach successfully captured the novel protein-protein interaction pattern that distinguishes this structure from the other 10 PDB structures of the same complex. The significant improvement in prediction accuracy demonstrates the model’s ability to correctly identify previously unseen ternary configurations despite never encountering this specific PPI pattern during training. Our approach precisely predicted the binding mode of the molecular glue and accurately positioned the key interface residues (Figure 2(e)), validating the model’s capacity to learn potential atomic-level interactions and apply them to novel structural arrangements.

Similar to the VHL:MG:CDO1 case, we observed substantial improvement when employing FK steering. As shown in Figure 4, FKSFold-Chai achieved a DockQ score of 0.590 with iRMSD = 1.813 Å, significantly outperforming Chai-1’s baseline model (DockQ = 0.076, iRMSD = 9.172 Å). This result demonstrates FKSFold’s generalization capability by correctly identifying novel PPI patterns not present in the training set.

**Figure 4:**
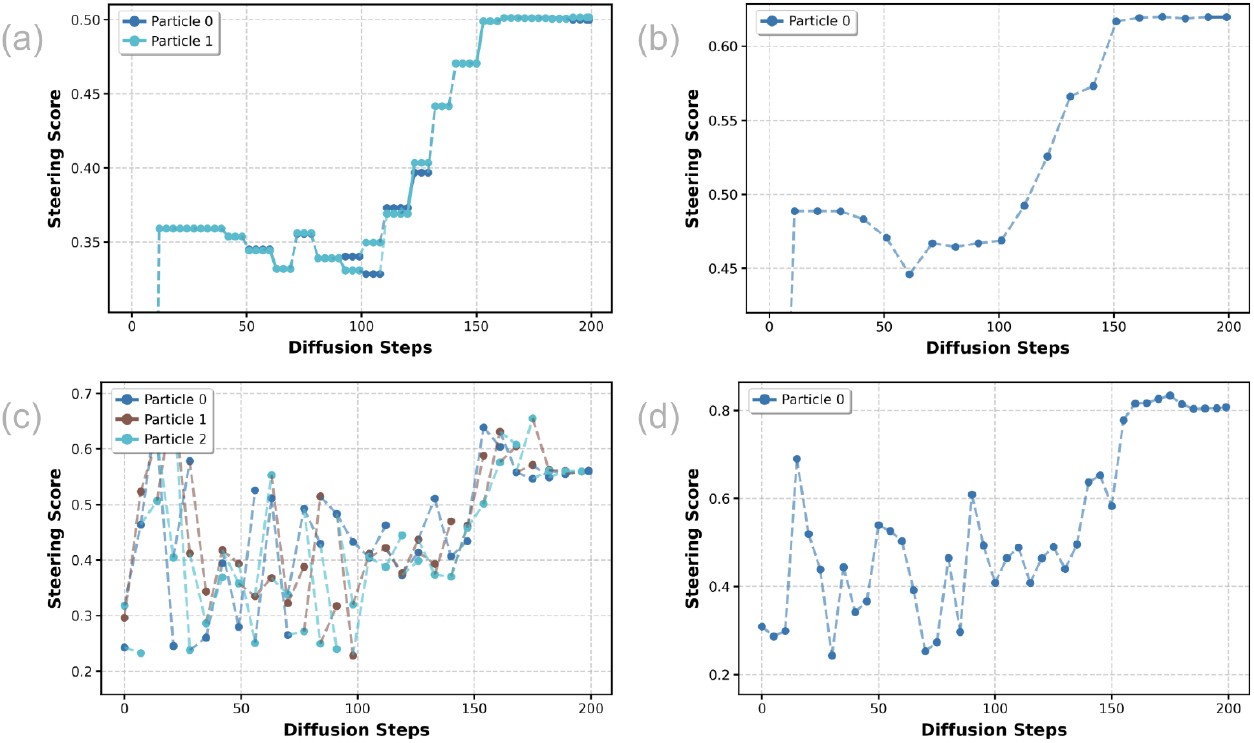
Internal diffusion score trajectories for the FKBP12:MG:mTOR-FRB, (a) Chai-1 with ipTM-guided steering and (b) Chai-1 default diffusion and (c) Boltz-2 with ipTM-guided steering and (d) Boltz-2 with default diffusion

**Figure 5:**
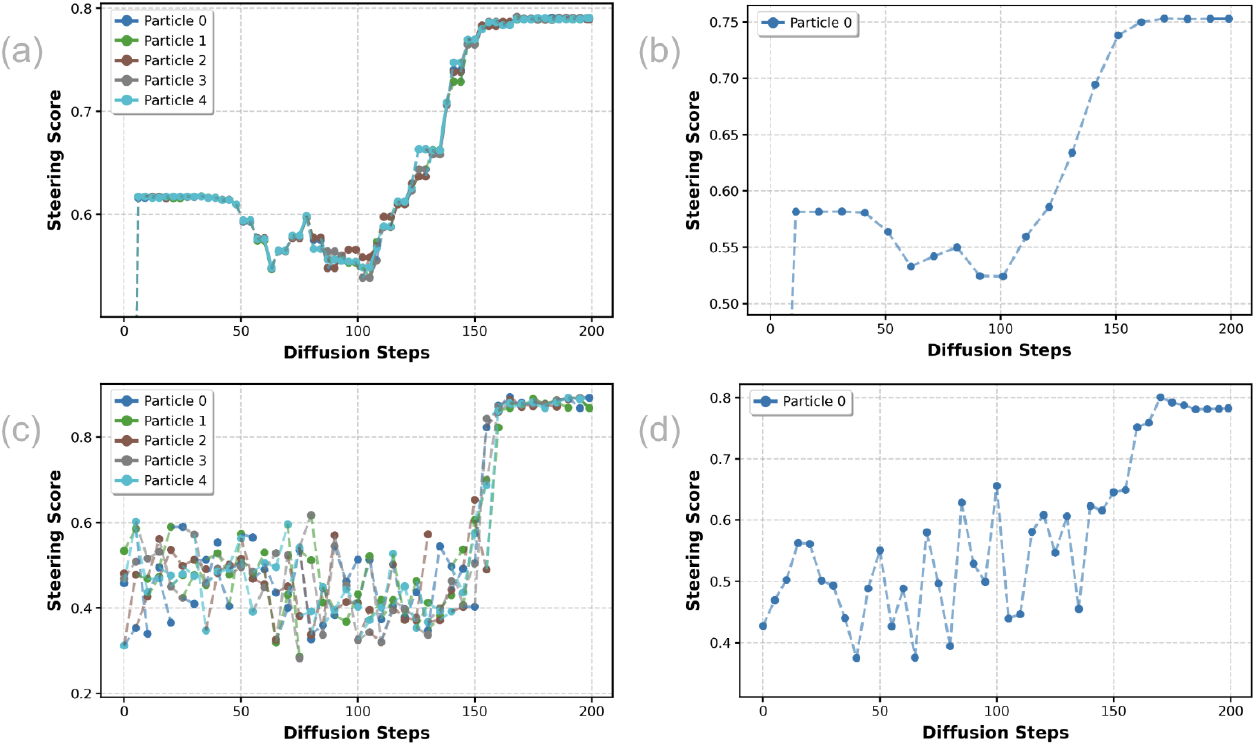
Internal diffusion score trajectories for the FKBP12:MG:BRD9, (a) Chai-1 with ipTM-guided steering and (b) Chai-1 default diffusion and (c) Boltz-2 with ipTM-guided steering and (d) Boltz-2 with default diffusion

This observation reveals a fundamental insight into the nature of diffusion-based structure prediction for molecular glue systems. The effectiveness of FK steering appears to be system-dependent, with different molecular glue complexes responding variably to the steering approach.

This system-dependent behavior manifests differently across model architectures. For Chai-1, we observe that the baseline model can occasionally achieve reasonable accuracy (DockQ = 0.555), with FK steering providing modest but consistent improvements. In contrast, Boltz-2 shows more pronounced baseline limitations but greater potential for improvement through steering, though this comes with increased variability due to scoring instabilities in its confidence head architecture.

#### FKBP12:MG:BRD9: Another Challenging Non-E3 Ligase Molecular Glue System

FKBP12:MG:BRD9 illustrates how molecular glues can mediate novel protein-protein interactions between FKBP12 and BRD9 to modulate epigenetic regulators, creating a complex interface geometry that was previously difficult to model. Figure 2(c) shows the comparison between FK steering prediction and experimental structure.

Our FK steering approach significantly improved prediction accuracy for this system, achieving a DockQ of 0.841 with iRMSD = 0.801 Å and fnat = 0.913, correctly identifying the binding mode of the molecular glue and the configuration of the interface (Figure 2(f)). This represents a substantial improvement over the baseline Chai performance (DockQ = 0.258, iRMSD = 2.569 Å, fnat = 0.250) and demonstrates consistent enhancement across all evaluation metrics.

### Analysis of Remaining Challenging Cases

For the four cases where FK steering showed improvements but did not achieve successful predictions, we observed several common challenges:

1. Flexible protein regions: CRBN:MG:VAV1-SH3c involves interfaces with highly flexible loop regions that undergo significant conformational changes upon binding
2. Large conformational sampling space: CRBN:MG:NEK7, FKBP12:MG:QDPR and KBTBD4:MG:HDAC1 complexes present greater conformational sampling challenges due to their larger protein sizes. NEK7 (283 residues), QDPR (230 residues) and HDAC1 (369) are significantly larger than proteins in successful cases, such as CDO1 (185 residues) and BRD9 (98 residues) when we used them as the inputs. This expanded conformational space makes comprehensive structural sampling more difficult, likely contributing to prediction failures.

Figure 6 illustrates these challenges using CRBN:MG:VAV1-SH3c as an example. This case highlights the difficulty of modeling flexible protein regions, specifically the RT-loop from VAV1 SH3c domain that engages in binding to CRBN. Conformational sampling of flexible long loops remains challenging. As shown in Figure 6(b), key residues D797 and R798 from the RT-loop in our FK steering prediction display significant positional deviations in both backbones and side chains compared to YDS-GlueFold prediction. Notably, the oxygen atom from the carboxylate group of D797, which normally forms a critical hydrogen bond with H357 from CRBN, is mispositioned. This hydrogen bond is essential for stable PPI formation between CRBN and VAV1-SH3c. The inherent flexibility of the VAV1 RT-loop increases prediction difficulty and likely contributes to the limitations of the FK steering method.

**Figure 6:**
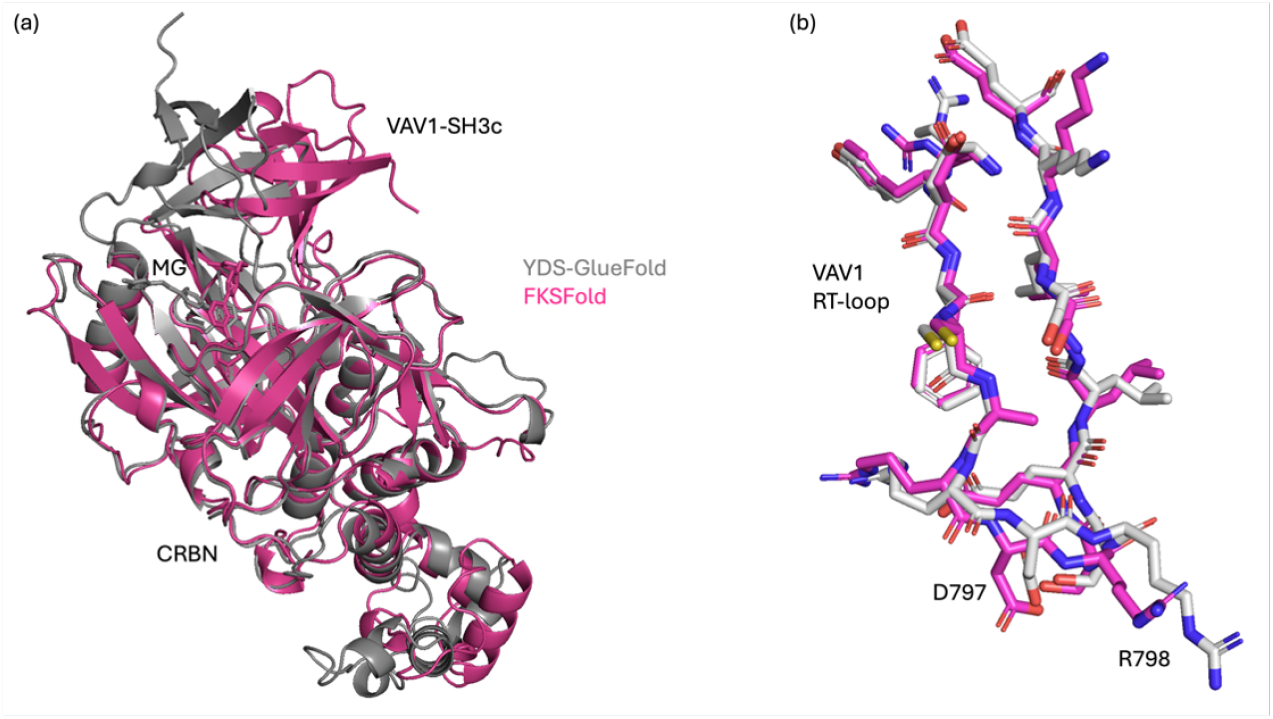
Analysis of remaining challenges in CRBN:MG:VAV1-SH3c. (a) The reference structure (YDS-GlueFold, gray) compared with FK steering prediction (red). (b) RT-loop of VAV1 showing conformational differences between reference structure (gray) and FK steering prediction (red), highlighting significant conformational changes in residues D797 and R798.

### Hyperparameter Sensitivity in FK Steering

Our FK steering implementation incorporates several critical hyperparameters that significantly influence the balance between exploration and exploitation during the diffusion process for ternary complex prediction. Comprehensive sensitivity analysis across our test cases reveals how these parameters affect prediction accuracy and provides insights for optimal parameter selection. Our systematic parameter sensitivity analysis provides crucial insights into how different configurations affect prediction accuracy across our test cases. The results in Table 5 demonstrate that FK steering performance is significantly influenced by four key parameters: particle numbers, diffusion path length (resampling interval), lambda weight for the temperature parameter, potential function type for FK steering activation.

**Table 5:** Hyperparameter sensitivity results of RMSD (Å) for VHL:MG:CDO1, FKBP12:MG:mTOR-FRB and FKBP12-MG-BRD9 (parameter sensitivity analysis of FK steering [particle numbers, diffusion path length, lambda weight, potential type])

| **Component** | **VHL:MG:**<br>**CDO1 (↓)** | **FKBP12:MG:**<br>**mTOR-FRB (↓)** | **FKBP12:MG:**<br>**BRD9 (↓)** |
| --- | --- | --- | --- |
| [2, 5, 10.0, vanilla] | 1.82 | 1.99 | 2.52 |
| [2, 5, 10.0, diff] | 1.60 | 1.47 | 2.51 |
| [2, 5, 10.0, max] | 1.76 | 1.99 | 2.51 |
| [2, 5, 20.0, vanilla] | 1.44 | 1.45 | 2.52 |
| [2, 5, 20.0, diff] | 1.74 | 1.46 | 2.53 |
| [2, 5, 20.0, max] | 1.80 | 2.02 | 2.51 |
| [2, 10, 10.0, vanilla] | 2.28 | 4.57 | 2.36 |
| [2, 10, 10.0, diff] | 3.16 | 4.52 | 2.30 |
| [2, 10, 10.0, max] | 2.12 | 4.53 | 2.38 |
| [2, 10, 20.0, vanilla] | 3.09 | 4.52 | 2.41 |
| [2, 10, 20.0, diff] | 2.13 | 4.56 | 2.32 |
| [2, 10, 20.0, max] | 2.19 | 4.52 | 2.38 |
| [5, 5, 10.0, vanilla] | 1.90 | 4.86 | 2.52 |
| [5, 5, 10.0, diff] | 1.83 | 4.56 | 2.54 |
| [5, 5, 10.0, max] | 1.45 | 4.93 | 2.14 |
| [5, 5, 20.0, vanilla] | 1.96 | 4.97 | 2.54 |
| [5, 5, 20.0, diff] | 1.76 | 1.71 | 2.38 |
| [5, 5, 20.0, max] | 1.42 | 4.92 | 2.14 |
| [5, 10, 10.0, vanilla] | 2.60 | 1.81 | 10.55 |
| [5, 10, 10.0, diff] | 4.17 | 1.78 | 10.61 |
| [5, 10, 10.0, max] | 3.16 | 1.80 | 10.58 |
| [5, 10, 20.0, vanilla] | 3.58 | 1.80 | 2.73 |
| [5, 10, 20.0, diff] | 2.77 | 4.56 | 2.29 |
| [5, 10, 20.0, max] | 3.49 | 1.79 | 10.74 |

The sensitivity varies considerably between different molecular systems. For VHL:MG:CDO1, all parameter settings yielded correct predictions with RMSD values consistently below 2.0 Å, suggesting this represents a relatively straightforward case where prediction accuracy remains robust across different parameter configurations. However, the other two cases demonstrated significant parameter sensitivity. In the FKBP12:MG:mTOR-FRB system, the parameter setting [2, 5, 10.0, vanilla] failed to produce accurate predictions with RMSD 1.99Å, while simply increasing the diffusion path length from 5 to 10 resulted in correct structural prediction with RMSD 4.57Å. Conversely, for FKBP12:MG:BRD9, decreasing the diffusion path length from 10 to 5 was necessary to achieve accurate results. These findings highlight the importance of parameter tuning for practical applications of our method, particularly for more challenging molecular glue systems where prediction accuracy is highly dependent on optimal parameter selection.

#### Lambda Weight

The lambda weight directly controls the concentration of sampling probability mass. Higher values of *λ* (e.g., 20.0) amplify differences between interface quality scores, strongly favoring particles with superior ipTM scores and potentially leading to less diverse sampling. Lower values (e.g., 10.0) flatten the probability distribution, allowing lower-scoring particles to survive resampling with higher probability, thus promoting exploration of alternative binding modes. Our ablation studies demonstrate that optimal *λ* values depend on the specific ternary system and sampling parameters. For VHL:MG:CDO1, a higher *λ* value of 20.0 with parameters [2, 5, 20.0, vanilla] yielded the best results (RMSD 1.44Å), while for FKBP12:MG:mTOR-FRB, a moderate *λ* value of 20.0 with parameters [2, 5, 20.0, vanilla] produced optimal predictions (RMSD 1.45Å).

#### Potential Type

The choice of potential type affects system convergence characteristics. The vanilla potential prioritizes absolute interface quality, while the difference potential rewards relative improvement, potentially preventing premature convergence. The maximum potential balances between historical success and current promise, preserving high-scoring trajectories while allowing exploration of newly promising pathways. Our experiments indicate that for VHL:MG:CDO1, the different potential types showed varying performance with diff potential achieving 1.60Å, vanilla potential 1.82Å, and max potential 1.76Årespectively. However, for FKBP12:MG:BRD9, the max potential showed improved performance (2.14Å) compared to vanilla (2.52Å) when using parameters [5, 5, 10.0].

#### Particle Numbers and Diffusion Path Length

The number of particles and diffusion path length parameters interact critically to determine sampling quality. For VHL:MG:CDO1, using 2 particles generally outperformed 5 particles, with the best configuration [2, 5, 20.0, vanilla] achieving 1.44Å. For FKBP12:MG:mTOR-FRB, increasing particles from 2 to 5 significantly improved performance in some cases, with [5, 5, 20.0, diff] achieving 1.71Åcompared to 1.45Åfor the 2-particle equivalent. The diffusion path length showed system-dependent effects: shorter paths (5 steps) generally performed better for VHL:MG:CDO1 and FKBP12:MG:mTOR-FRB, while FKBP12:MG:BRD9 showed complex behavior with some configurations performing poorly at longer path lengths (e.g., 10.55Åfor [5, 10, 10.0, vanilla]).

These hyperparameters interact in complex ways, necessitating careful tuning for optimal performance on specific molecular glue systems. The parameter combination [2, 5, 20.0, vanilla] yielded the best overall performance for both VHL:MG:CDO1 (1.44Å) and FKBP12:MG:mTOR-FRB (1.45Å), though system-specific optimization remains important for challenging ternary complexes.

## Conclusion

This paper has presented technical details of our early experiments with Feynman-Kac steering for molecular glue-induced ternary complex prediction—work that predated our successful YDS-GlueFold model. While FK steering achieved only partial success, correctly predicting 3 out of the 8 test cases that were later solved by YDS-GlueFold, it represented an important developmental step in our technical journey. The approach demonstrated the potential of using stochastic control theory to guide diffusion processes toward high-quality protein-protein interfaces in ternary complexes, with ipTM scores serving as a biologically relevant reward function. This alignment between our computational approach and the underlying mechanism of molecular glues provided a sound conceptual foundation, despite technical limitations in our early implementation.

The partial successes in cases with well-defined interfaces demonstrated the promise of this approach, while the failures in more challenging cases highlighted the need for more sophisticated methods. By documenting this early approach after having already reported the successful results of YDS-GlueFold, we aim to provide a more complete picture of the technological evolution in this challenging area of computational chemistry. The identified limitations of FK steering illustrate the iterative nature of method development in the complex field of protein-ligand-protein interaction modeling.

The technical details presented here may be of value to other researchers working on related challenges, potentially informing alternative approaches or hybrid methods. As the field continues to advance, some of the concepts explored in our FK steering experiments may find new applications or inspire novel methodological innovations in computational chemistry and structural biology.

### Future Directions for FK Steering Engineering

Our exploration of FK steering reveals multiple promising avenues for further development and optimization. Recent work, Boltz-1x, ^20^ has already improved physical plausibility of structure prediction by using force field-inspired potentials to steer diffusion. Extending from these advances, numerous variants can be explored to further enhance the effectiveness and applicability of FK steering in molecular structure prediction:

#### Advanced Potential Function Design

Generally, the steering objective must be aligned with the task goal. In this work, we utilized ipTM as the scoring criterion because we prioritize the generation quality of interface regions. Beyond the potential function variants explored in this work, more sophisticated designs could incorporate multi-objective potentials that simultaneously optimize interface quality, physical plausibility, and ligand binding interactions with appropriate weighting schemes, or hierarchical potentials that operate at different structural resolution levels throughout the diffusion process. Physical potentials *U* (*x*) are functions defined on structural coordinates *x*, which are applicable even to inference-only models such as Chai-1.

For models with fully open-sourced weights, such as Boltz-2 or Protenix, access to model gradients enables more flexible model modifications, as demonstrated in Boltz Design,^27^ potentially supporting more advanced potential function implementations. Finally, we observed that the confidence head scoring of certain open-source models exhibits instability. Training specialized scoring models may represent a viable alternative approach.

#### Embedding Selection Strategies

In AlphaFold3, the PairFormer replaces the Evoformer as the representation learner. It consists of a stack of attention with pair-bias, triangle operations, gated MLPs, layer norms, and recycling. None of these components include inference-time randomness by default (no dropout, no stochastic depth at test), so identical inputs produce identical outputs.

We observed the substantial influence of PairFormer embeddings on the diffusion process, including MSA quality and whether PairFormer embeddings are generated under no-MSA/no-template conditions. Methods for noise filtering in latent diffusion initialization^17^ may provide valuable insights. Introducing stochasticity in the PairFormer may help identify superior conformations.

#### Alternative Particle Selection Strategies

Rather than selecting single particles for continuous diffusion sampling, more robust approaches could include importance mixing that combines particles from different rounds of sampling to prevent premature convergence, or staged filtering that applies increasingly stringent selection criteria as diffusion progresses.

Additionally, our diffusion filtering satisfies a decision tree data structure. Search efficiency can be improved through algorithms such as depth-first search (DFS), breadth-first search (BFS), or backtracking.^28^

#### Steering-Aware Training

Alternative perspectives on diffusion models exist. Taking DDPM as an example, DDPM can be conceptualized as an Autoregressive VAE. Theoretically, Chain-of-Thought (CoT) for large language models increases total computational resources for problem-solving through multi-token outputs, thereby improving LLM performance. Similarly, diffusion models could enhance performance by increasing diffusion steps. Unfortunately, we observed that brute-force increases in diffusion steps do not contribute to model performance, possibly because diffusion and noise scheduler training do not account for steering factors.

A subsequent consideration involves incorporating partial diffusion and variable diffusion step steering into the training process, which would significantly improve steering performance.^29^

## TOC Graphic

Some journals require a graphical entry for the Table of Contents. This should be laid out “print ready” so that the sizing of the text is correct. Inside the tocentry environment, the font used is Helvetica 8 pt, as required by *Journal of the American Chemical Society*.

The surrounding frame is 9 cm by 3.5 cm, which is the maximum permitted for *Journal of the American Chemical Society* graphical table of

content entries. The box will not resize if the content is too big: instead it will overflow the edge of the box.

This box and the associated title will always be printed on a separate page at the end of the document.

